# ScProteoAgent enables natural-language-driven single-cell proteomics analysis and interpretation

**DOI:** 10.64898/2026.09.24.754011

**Authors:** Runwen Hu, Keyan Ding, Wenwen Wang, Zhihui Zhu, Peilin Chen, Shiqi Wang, Yu Wang

## Abstract

Single-cell proteomics requires computational choices to remain aligned with experimental design as research questions evolve. Here we present ScProteoAgent, which translates natural-language requests into domain-specific calculations and preserves research intent, analysis design, matrix-processing history and statistical outputs as reusable analytical state. Follow-up requests reuse applicable inputs and initiate new calculations when contrasts or experimental units change. We assembled a benchmark comprising 55 tasks and 33 reference conclusions from 11 published studies, combining progressive questions within studies with complementary biological and analytical settings. In a retrospective comparison of 88 archived outputs from eight systems, ScProteoAgent achieved the highest mean score of 93.67 under a common six-dimensional rule-based evaluation. A HeLa migration case connected phenotype composition with proteomic clusters, candidate differences and original-observation support. In liver zonation, a researcher-guided follow-up reused a saved matrix to test mouse-level spatial contrasts and compare full-matrix and observed-only results. Applications in brain development, sample preservation and hematopoiesis illustrated how interpretation depends on processing choices and experimental units. ScProteoAgent connects research questions with statistical results and their analytical context, enabling existing analyses to support more specific biological investigations.

## Introduction

Single-cell proteomics provides direct access to protein abundance and coordinated variation, offering a distinctive view of cellular heterogeneity^1, 2, 3^. Advances in measurement depth, throughput and the preservation of spatial and phenotypic information have expanded research questions from distinguishing cell types to resolving continuous states, interpreting environmental responses and understanding functional organization within tissues^4, 5, 6^. Cells of the same type can express different protein programs, and similar proteomic states can encompass different phenotypes. Turning these measurements into biological understanding therefore requires protein changes to be interpreted in relation to cell origin, experimental treatment and tissue location, with individual candidates considered alongside population-wide patterns.

Identification and quantification software provides the data foundation for this work. DIA-NN, Spectronaut and MaxQuant support processing from raw mass spectrometry data to peptide or protein quantification^7, 8, 9^, while PEAKS combines *de novo* sequencing with database searching to obtain peptide sequence evidence^10^. Once a matrix is available, downstream analysis must be organized around the research question. Perseus offers established statistical and visualization capabilities^11^, and frameworks such as scp and scplainer support standardized data organization, flexible processing and interpretable modeling^12, 13^. Researchers also routinely develop customized R and Python scripts. These methods provide a rich computational foundation; applying them to a particular study requires sustained connections among scientific questions, data properties, tool parameters and interpretation.

These connections have specific implications for single-cell proteomics. Filtering, missing-value handling, normalization and batch correction interact, influencing the retained protein set, the scale of estimated differences and their interpretation^14, 15^. Imputation, for example, enables subsequent calculations on sparse matrices but changes the observational support underlying candidate effects. Processing choices must therefore accommodate both the data and the biological difference to be estimated. Cell size, protein detection depth and sample preservation can also influence measured abundance patterns. Cells, donors and tissue sampling locations represent different levels of observation: variation among cells from one donor and changes reproducible across donors answer different questions. Statistical inference in hierarchical sampling must reflect the experimental unit^13, 16, 17^. Matrix state, target and reference groups, experimental units and technical factors consequently need to remain explicit from task definition through computation and interpretation.

Scientific analysis also develops as results emerge. A population enriched for migrating cells prompts questions about its composition and molecular candidates; differences between spatial endpoints can motivate a direct test of the intervening region. Such questions may require revised contrasts, a different aggregation unit or sensitivity analysis of processing choices. Preserving comparison definitions, statistical results and intermediate matrices allows completed work to become the starting point for the next analysis. Research object and workflow provenance approaches already provide machine-readable representations of inputs, outputs and their relationships^18, 19^. In interactive analysis, these records must also inform executable choices about which inputs remain applicable and which results need to be recalculated. Agents that interpret natural-language requests and coordinate tools provide a new interface for this movement between questions, data and interpretation.

Biomedical agents have demonstrated task planning, tool use and iterative exploration. Biomni addresses a broad range of biomedical research tasks^20^, while CellVoyager and SpatialAgent illustrate agent-based investigation of cellular data and spatial biology^21, 22^. PROTEUS combines language models with specialized tools for exploratory proteomics and organizes multiomics research through structured relationships among research directions, tools and results^23, 24^. SciToolAgent uses a scientific tool knowledge graph to support tool selection and execution^25^. These approaches establish tool coordination as a practical interface to scientific computation. In single-cell proteomics, that coordination must also preserve the experimental meaning of each calculation. Questions need defined groups, directions and experimental units; matrices need processing histories; and subsequent analyses need access to the conditions underlying earlier results. We focus on carrying this design and processing context from a research request through computation to a follow-up question.

Here, we introduce ScProteoAgent, which organizes domain-specific computation around scientific questions, preserves designs and results as reusable analytical state, and supports researchers in refining their questions. Three connected principles guide the system. *Question-centric analysis* translates research requests into analytical objects, contrasts, experimental units and tool parameters. *Persistent analytical state* connects research intent, analysis design, data state and result state while preserving their dependencies. *Continuable scientific analysis* allows follow-up questions to reuse applicable products and trigger the new computations they require. Building on this framework, we develop a benchmark that combines progressive questions within studies with complementary challenges across studies, apply a common rule-based evaluation to eight systems, and use migration and liver spatial studies to demonstrate complementary roles in phenotype interpretation and question refinement. Together, the system, benchmark and applications establish a practical route from natural-language research ideas to specialized proteomic analysis.

## Results

### Connecting research tasks with domain computation

ScProteoAgent receives a protein quantification matrix, sample metadata and a natural-language request, and produces a report, statistical tables, figures and saved analytical outputs (Fig. 1a). These inputs define observations, experimental context and research intent. Matrix columns are linked to sample identifiers and separated from protein annotations. Membrane-state comparisons require intact and permeable labels together with storage conditions and cell types; spatial comparisons require zone labels and, when available, identifiers for biological individuals.

**Fig. 1.**
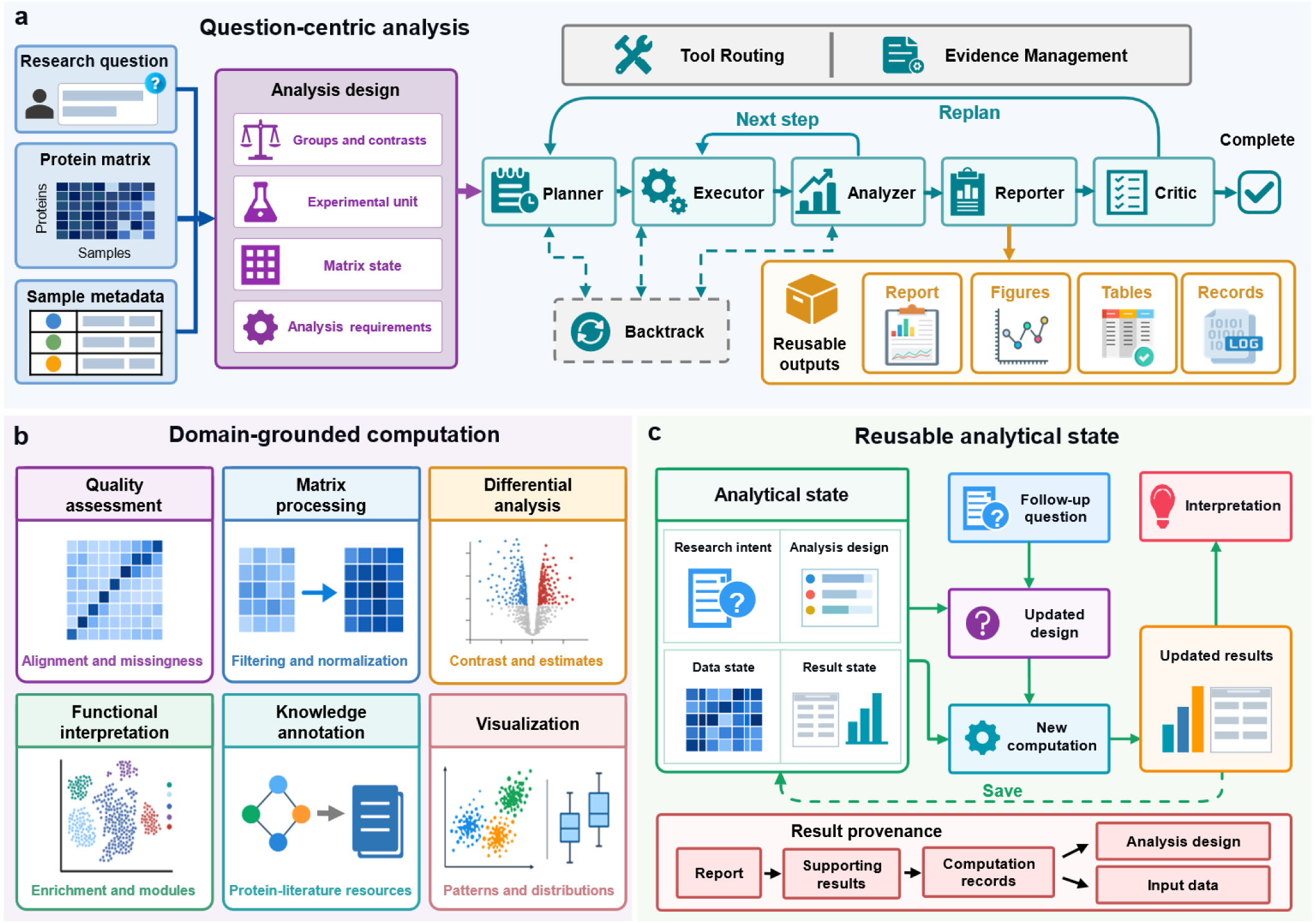
ScProteoAgent integrates question-centric analysis, domain-grounded computation, and reusable analytical state. **a**, Question-centric analysis workflow. A research question, protein quantification matrix, and sample metadata inform the analysis design, including groups and contrasts, experimental units, matrix state, and analysis requirements. Planner, Executor, Analyzer, Reporter, and Critic coordinate analysis and reporting, with replanning and backtracking supporting workflow revision. Tool routing and evidence management support the workflow, which produces reports, figures, tables, and execution records.**b,** Domain-grounded computational capabilities spanning quality assessment, matrix processing, differential analysis, functional interpretation, knowledge annotation, and visualization.**c,** Reusable analytical state links research intent, analysis design, data state, and result state. A follow-up question informs an updated design and new computation using applicable saved state. Updated results support interpretation and are saved for subsequent analysis. Result provenance connects reported findings to supporting results, computation records, analysis design, and input data.

Task representation translates research language into computational conditions. Grouping variables, target and reference groups, matrix state and experimental units define the analysis design. The design retains information sources, distinguishing supplied metadata from conditions requiring confirmation. For example, comparing the midlobular zone with the average of the two endpoints gives weights of 1 for the midlobular zone and −0.5 for each endpoint. Adding mouse identifiers makes aggregation by mouse and zone necessary before the paired comparison. The same research question therefore determines both the contrast and the level of analysis.

Six node types coordinate execution: Planner, Executor, Analyzer, Reporter, Critic and Backtrack. Planning, execution and analysis form the main computational path, followed by report assembly and review. The Analyzer can return to the Executor for the next step, whereas the Critic can complete the task or initiate replanning. Conditional recovery through Backtrack restores execution at an appropriate node or returns the task to planning. Fig. 1a summarizes the principal execution and recovery paths. Tool routing and evidence management support coordination across these steps.

Analytical state has four connected components. Research intent records the question and task requirements. Analysis design specifies groups, contrast directions and experimental units. Data state identifies the applicable matrix and processing history. Result state connects statistical tables, summaries and figures with their dependencies. Shared objects carry this information within a run; saved design files and artifact indexes make it accessible to subsequent runs. The liver continuation retains the parent question and processed matrix while adding mouse mapping, contrast weights and the resulting unit-level statistics. This connection allows a later computation to use the intended data under an explicit design. Evidence records connect reported findings to supporting statistical results and source artifacts. Saved designs and matrix-processing records locate each result within its contrast, experimental unit and input data (Fig. 1c).

Domain tools support six complementary capabilities (Fig. 1b). Quality assessment checks sample matching, protein coverage and missingness. Matrix processing filters, transforms and imputes data according to input state. Differential analysis returns effect estimates and tests for defined contrasts. Functional interpretation connects proteins to saved modules and enrichment results. Knowledge interfaces support annotation of proteins, pathways and interactions, while visualization presents sample structure and statistical results. Tools return parameters, analytical objects and output locations alongside summaries. Full matrices and statistical tables remain in the analysis package; the model reads compact summaries containing contrasts, representative findings and execution status.

The continuation interface supplies the full parent task, saved design and artifact index to a new task (Fig. 1c). Parent outputs remain read-only, and new calculations are saved separately. A new contrast can reuse an applicable matrix, while a changed experimental unit requires reaggregation and new statistics. Revised processing conditions similarly define a separate analytical branch. Reuse and recalculation therefore occur together: the saved state identifies the input and its conditions, and the updated design specifies the calculation. In the liver case, this interface connects an existing spatial analysis to direct mouse-level tests without regenerating its data foundation.

Reports link interpretations to structured comparisons and numerical results. Researchers can inspect the underlying tables or reuse their associated matrices and designs. The applications below illustrate how these connections support phenotype interpretation, spatial comparisons and analyses of experimental context.

### A benchmark of progressive and complementary questions

We constructed a research-oriented benchmark to examine how agents address connected tasks under different data conditions (Fig. 2a and Supplementary Fig. 1). It combines protein quantification matrices and sample information from 11 published studies, encompassing 5,262 measured objects, including single cells, tissue sampling units and low-input samples. Observation types and replicate structures are recorded for each study, distinguishing matrix size from the experimental unit. Study names describe the biological object and question, such as THP-1 inflammatory responses, liver zonation and pancreatic tumor T-cell states.

**Fig. 2.**
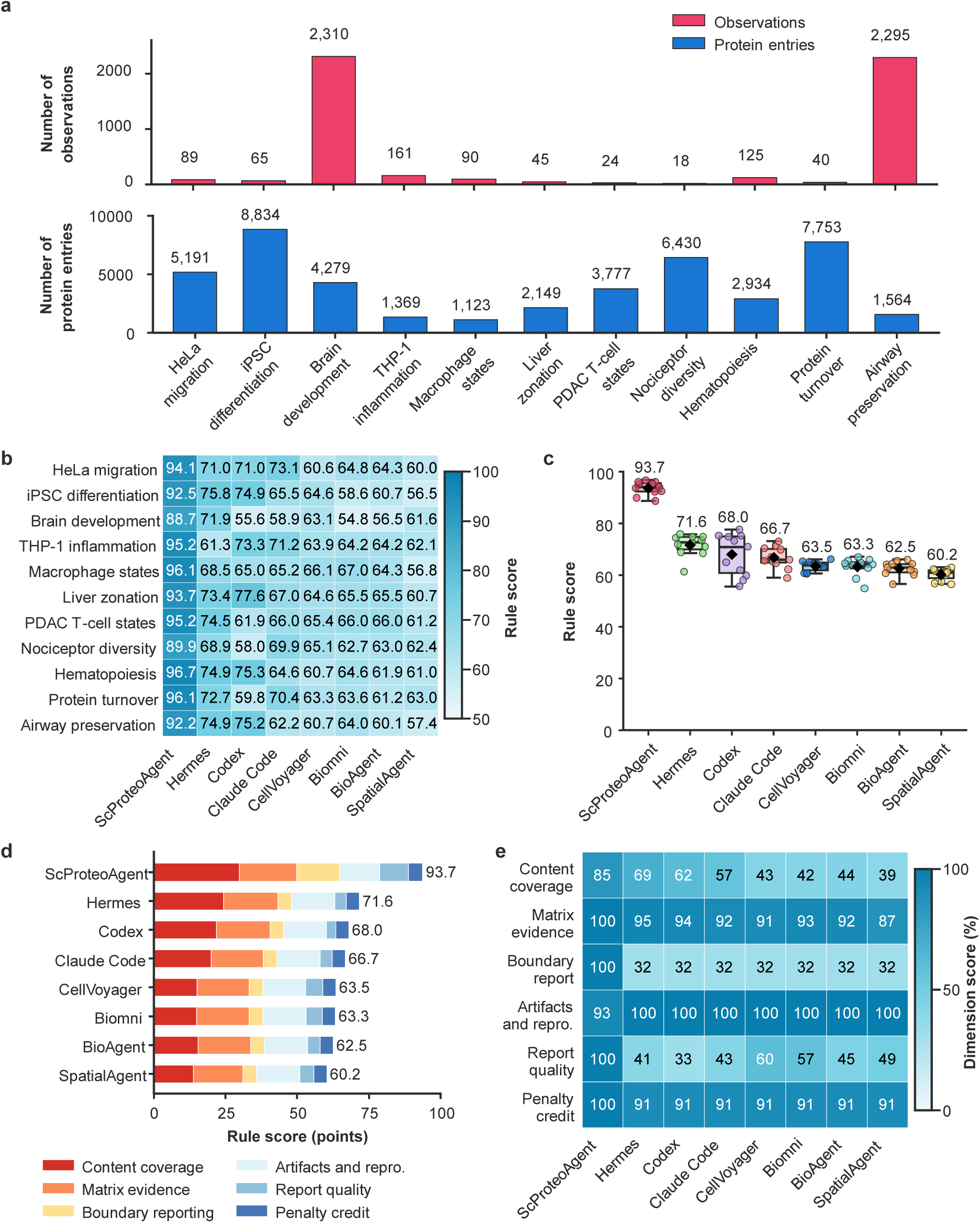
A research-oriented benchmark and cross-system rule-based evaluation. **a**, Observation counts and input protein-row counts for 11 studies, displayed on separate linear axes starting at zero. The 5,262 observations include single cells, spatial sampling units and low-input or pooled samples. Protein rows are not summed across studies as unique proteins. **b,** The 88 rule totals for eight systems across 11 studies. Scores range from 0 to 100 by definition; the displayed color scale spans 50–100 and contains all observed scores. **c,** Study-level scores, with 11 points per system. Boxes show the 25th–75th percentiles, center lines the median and whiskers the most extreme observations within 1.5 interquartile ranges. Black filled diamonds show means, also displayed numerically to one decimal place. **d,** Mean contributions of the six dimensions across studies. Segment sums equal the mean composite score. Maximum scores are 35 for research content, 20 for matrix evidence, 15 for interpretive boundaries, 15 for artifacts and reproducibility records, 10 for report quality and 5 for the score retained after penalties. **e,** Mean dimension scores divided by their respective maxima, expressed as percentages rounded to the nearest integer without percent signs in the cells. These are proportions of available dimension points, not contributions to the composite total. All panels use the same deterministic rule evaluation, without historical language-model adjustments or additional normalization. Paired comparisons and fixed checks are shown in Supplementary Figs. 2 and 3.

The studies were selected for complementary scientific questions. Migration and differentiation connect phenotypes and cell states with protein programs. Membrane integrity introduces storage conditions and cell composition. Spatial zonation combines positional relationships with repeated sampling. Neuronal subtypes and immune states provide settings for comparisons across biological contexts. The studies also differ in missingness and input processing state. These differences allow the benchmark to examine how domain information shapes an analytical path, rather than testing only one fixed sequence of calculations.

From the questions and applications in the source publications, we manually organized five related analytical requirements and three reference conclusions per study, yielding 55 tasks and 33 reference conclusions. The five requirements were recorded together in each study’s task request. Their progression connects grouping and state analysis, candidate identification and integrated biological interpretation; it describes the structure of the benchmark rather than five separately administered interaction rounds. Questions range from population composition and mixed populations to experimental context, spatial relationships and functional organization. Each study retains one question structure within a complementary biological setting.

Each benchmark instance contains a matrix, metadata, task text, reference conclusions and evaluation criteria. The first three define the analytical problem; the source study provides the basis for the reference conclusions and scoring requirements. This correspondence allows evaluation to address concrete research content: whether key comparisons are represented, quantitative findings are expressed and conclusions are connected to evidence. Preserving all five components makes task construction and evaluation inspectable and reusable.

A single deterministic rule function evaluates reports and archived analysis packages from eight systems across the 11 studies. Its six components cover research content, matrix evidence, interpretive boundaries, artifacts and reproducibility records, Chinese report quality, and the score retained after penalties, with maxima of 35, 20, 15, 15, 10 and 5 points. Requirements derived from the source publications provide content references. The program applies fixed checks for reference terms, key contrasts, numerical expressions, report structure and artifact records, then aggregates them with fixed weights. The same function and input conventions are applied to all 88 outputs, retaining the component-level scoring basis. This provides an explicit and repeatable computation for the comparison.

Progressive tasks within studies and complementary conditions across studies jointly define the benchmark’s value. The former examine how a system connects earlier results to subsequent questions within a study; the latter examine its applicability across experimental structures. The benchmark therefore serves both as a shared task collection for system comparison and as a resource for locating analytical gaps and designing further research tests.

### Rule-based comparison of archived analysis outputs

We retrospectively compared 88 archived analysis outputs from ScProteoAgent, Biomni, SpatialAgent, BioAgent, CellVoyager, Codex, Claude Code and Hermes across 11 studies. GPT-5.5 was the base model for ScProteoAgent in this comparison. Reports and archived products were scored using the same six-dimensional rule function, with studies as the units of aggregation. The archived collection includes reference-linked evaluation material and report content; its relationship to model inputs is only partly documented (Methods). The comparison characterizes archived outputs under their historical generation conditions. Study-level scores and component checks are shown in Fig. 2b–e and Supplementary Figs. 2 and 3.

**Fig. 3.**
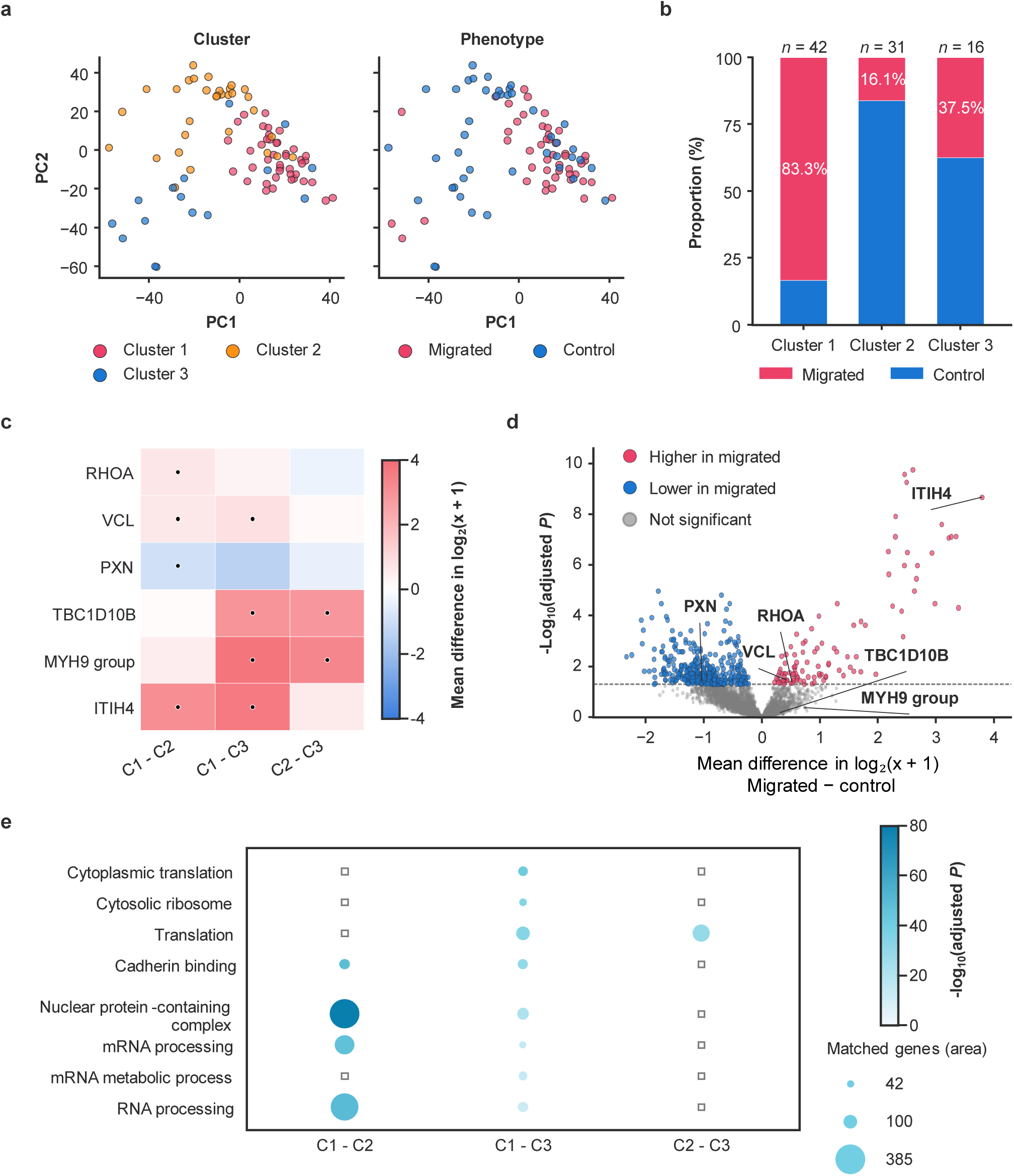
Connecting migration phenotypes, population composition and protein candidates. **a**, Identical principal-component coordinates for 89 HeLa cells, colored by source-provided cluster labels or migration labels and displayed with shared coordinate ranges. PC1 and PC2 explain 12.05% and 10.49% of the variance in the standardized protein matrix, respectively. Cluster labels are analysis inputs. **b,** Clusters C1, C2 and C3 contain 42, 31 and 16 cells, including 35, 5 and 6 migrated cells, respectively. Migrated fractions are 83.3%, 16.1% and 37.5%. Red and blue indicate migrated and control cells. The migration-composition odds ratio for C1 relative to C2 is 26.0 (95% confidence interval, 7.4–91.2). **c,** Mean differences for six fixed candidates across three cluster contrasts on the log₂(*x* + 1) abundance scale. Positive values indicate higher abundance in the first-named cluster. The color scale is centered at zero. Black dots indicate within-contrast Benjamini–Hochberg (BH) *q* < 0.05 across 4,688 proteins. **d,** Offline comparison of 46 migrated and 43 control cells in the saved matrix. Each point is a protein. The horizontal axis is migrated-minus-control mean log₂(*x* + 1) abundance; the vertical axis is −log₁₀ adjusted *P*. Two-sided Welch *t*-tests and one 4,688-protein BH family identify 513 *q* < 0.05 proteins, comprising 104 higher and 409 lower in migrated cells. The dashed line indicates *q* = 0.05, with no additional effect cutoff. Labeled proteins correspond to the fixed candidates in c. **e,** Saved GO results using the historical 19,591-gene background. Rows 1–4 (cytoplasmic translation, cytosolic ribosome, translation and cadherin binding) concern genes higher in the first-named cluster; rows 5–8 concern genes lower in that cluster. Terms were fixed as the four smallest adjusted *P* values per direction in C1 versus C3. Area encodes matched member genes; color encodes −log₁₀ adjusted *P*. Thirteen of 24 positions have saved results; open squares denote absence from the saved display tables, not nonsignificance. Experimental-background results (4,227 genes) are summarized in the text. TBC1D10B observation support is shown in Supplementary Fig. 4.

ScProteoAgent achieved the highest mean composite score, 93.67, with a range of 88.70–96.70 and a median of 94.10. Mean scores were 71.62 for Hermes, 67.96 for Codex, 66.73 for Claude Code, 63.46 for CellVoyager, 63.25 for Biomni, 62.52 for BioAgent and 60.25 for SpatialAgent. ScProteoAgent outscored every comparator in all 11 studies and exceeded the highest-scoring comparator, Hermes, by an average of 22.05 points. The study-level results show that this composite advantage spans different biological topics and data conditions (Fig. 2b,c).

The component scores identify the sources of this advantage. For research content coverage, ScProteoAgent averaged 29.76 of 35 points, compared with 24.25 for Hermes, 21.69 for Codex, 19.82 for Claude Code and 13.70–15.34 for the remaining systems. This component combines reference content, key findings and core contrasts, indicating that ScProteoAgent organizes a larger fraction of task-relevant information into its outputs. Matrix evidence reached 20.00 points, compared with 17.36–19.00 for the comparators, reflecting the connection among quantitative expressions, candidate evidence and access to results.

Interpretation and report organization provided another distinctive contribution. ScProteoAgent scored 15.00 for interpretive boundary reporting and 10.00 for Chinese report quality, compared with 4.73 and 3.27–6.00, respectively, for the comparators. These checks describe how matrix results, functional annotations and the scope of conclusions are organized in the report. For artifacts and reproducibility records, ScProteoAgent scored 13.91, whereas all comparators scored 15.00, indicating that the archived packages generally provided extensive structured records. The composite differences arose mainly from research-content checks and the required organization of evidence and report text.

The six-dimensional profile shows where the archived outputs differed: ScProteoAgent covered more reference-linked research content and fulfilled more evidence-reporting requirements, while comparator packages scored higher for artifact and run-state records. Presenting every study and dimension retains this distinction alongside the composite score.

### Connecting migration phenotypes to protein candidates

Cells from the same source can differ in motility. The PiSPA study combined a HeLa scratch-migration experiment with single-cell proteomic measurements to examine molecular differences between migrated and control cells^26^. In this setting, researchers need to identify populations associated with migration and connect them to proteins and functions. We analyzed the quantitative matrix for 89 cells together with migration labels and three cluster labels supplied by the source study, asking ScProteoAgent to organize state comparisons, identify candidates and interpret population heterogeneity (Fig. 3; Supplementary Fig. 4).

Population composition gives the proteomic states a phenotypic interpretation (Fig. 3a,b). Migrated cells constituted 83.3% of Cluster 1 (35/42), compared with 16.1% of Cluster 2 (5/31) and 37.5% of Cluster 3 (6/16). Cluster 1 therefore represents a proteomic state enriched for migrated cells, whereas the other two clusters are predominantly control cells but also contain migrated cells. This correspondence links subsequent cluster comparisons to migration while preserving heterogeneity within the same phenotype.

To connect migration labels directly to protein changes, an offline comparison of the saved matrix provided the effect distribution for 4,688 proteins, of which 513 passed adjustment within a single Benjamini–Hochberg (BH) family (Fig. 3d). This complements the composition analysis by placing label-associated changes alongside cluster heterogeneity. ScProteoAgent also generated differential tables for the three cluster contrasts, enabling candidate distributions to be interpreted with population composition (Fig. 3c). For TBC1D10B, the mean difference on the log₂(*x* + 1) scale was 2.93 for Cluster 1 relative to Cluster 3 (BH-adjusted *P* = 2.84 × 10⁻⁵) and 2.83 for Cluster 2 relative to Cluster 3 (2.24 × 10⁻⁵), but only 0.10 for Cluster 1 relative to Cluster 2 (0.866). In the processed matrix, TBC1D10B distinguished Cluster 3 from both other clusters, rather than separating the migration labels directly. Matching each value to the pre-imputation matrix confirmed that 31 of 89 values were imputed, including 14 of 16 values in Cluster 3, which had only two original observations (Supplementary Fig. 4). We therefore treated TBC1D10B as an imputation-sensitive, cluster-associated candidate for targeted measurement. The two observed C3 values provide limited support for establishing a cluster-specific abundance difference.

Functional enrichment connected the cluster contrasts to translation, adhesion and RNA processing (Fig. 3e). The figure preserves eight GO terms selected from the saved C1–C3 results. To examine dependence on the reference population, we repeated the same six directional queries using the 4,227 GO-annotated genes represented among the tested proteins. All eight C1–C3 terms retained support after this background restriction: translation and cadherin-binding terms were enriched among genes higher in C1, whereas RNA-processing and nuclear-complex terms were enriched among genes lower in C1. Cadherin binding also remained supported among genes higher in C1 than C2. Across the 24 fixed term–contrast positions, 15 were significant under both backgrounds, five only under the historical background and four under neither (*q* ≤ 0.05). The four C2–C3 higher-abundance terms and the C1–C2 translation term lost support with the experimental background. These results concentrate the functional interpretation on the supported cluster contrasts while preserving the distinction between cluster identity and migration phenotype.

Together, phenotype composition and protein contrasts distinguish migration-associated changes from differences between mixed proteomic populations. Observation records further identify which candidate differences require additional measurements in a specific population.

### Refining spatial questions through mouse-level comparisons

The liver lobule is functionally organized along the portal-to-central axis. The source deep visual proteomics (DVP) study combined tissue imaging, microdissection and mass spectrometry to examine hepatocyte proteomes at different positions^5^. Beyond recovering differences between the endpoints, this setting raises a more specific question: does the intervening midlobular zone have molecular features that warrant direct examination? Using quantitative data and labels for the three zones, we asked ScProteoAgent to organize zonal comparisons, regional candidates and metabolic interpretation (Fig. 4; Supplementary Fig. 5).

**Fig. 4.**
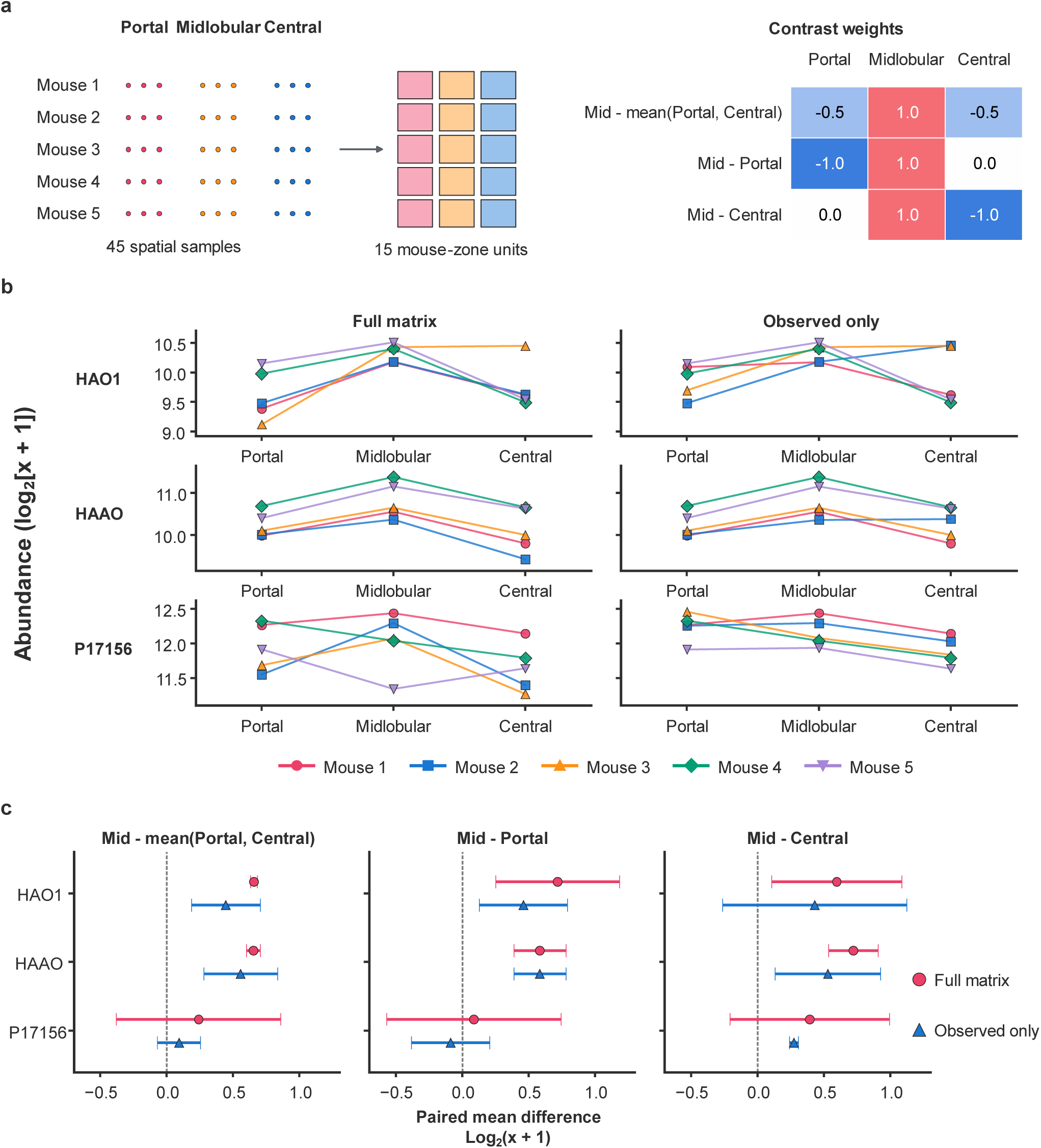
Refining liver zonation questions through experimental units and direct contrasts. **a**, Forty-five spatial sampling sites from five mice, with three zones per mouse and three sites per zone. Each site is a morphological pool of five hepatocytes. Within-zone aggregation produces 15 mouse-by-zone units. The weight matrix lists portal, midlobular and central coefficients for midlobular minus the endpoint mean and midlobular minus each endpoint. **b,** Aggregated HAO1, HAAO and P17156 abundances by mouse and zone. Full-matrix and observed-only branches are aggregated on the same log₂(*x* + 1) scale and share the vertical scale within each candidate. Each line represents one mouse, with consistent colors and shapes across the six facets. **c,** Eighteen paired mean differences on the log₂(*x* + 1) scale and unadjusted 95% *t* confidence intervals for three candidates, three contrasts and two branches, using five mice as paired units. The intervals are not corrected for multiple comparisons. Positive values indicate a higher midlobular value. Red circles and blue triangles represent full-matrix and observed-only estimates. Each branch is adjusted separately within contrasts and across the joint family of three contrasts using Benjamini–Hochberg correction; each adjusted result is interpreted within its own testing family. Candidate observation support, full-matrix support and both *q* values are provided in Supplementary Fig. 5.

The initial analysis established relationships among the zones. At the initial filtering thresholds, 154 proteins differed between the central and portal zones, 125 between the central and midlobular zones, and none between the midlobular and portal zones. The first two contrasts shared 57 proteins with higher abundance in the central zone, providing convergent regional evidence. The midlobular zone was already part of the initial task. These results sharpened the follow-up question: after accounting for repeated sampling within a mouse, is the midlobular deviation from the mean of both endpoints different from its comparisons with each endpoint separately?

In the follow-up, the researcher specified mouse units and three spatial comparisons, and the system updated the analysis design accordingly. The saved matrix and research context provided the starting point, while mouse identifiers and contrast weights defined the necessary reaggregation and testing. The dataset comprised 45 spatial observations from five mice, with three zones per mouse and three sampling sites per zone. The continuation reused the saved matrix with its recorded processing state, aggregated within each mouse and zone to obtain 15 mouse-by-zone combinations, and used mice as paired units. The contrasts were midlobular minus the mean of portal and central zones, midlobular minus portal, and midlobular minus central. These definitions yielded corresponding effects, confidence intervals and adjusted results, turning a broad midlobular question into explicit statistical comparisons (Fig. 4a).

In the full-matrix branch, which includes imputed values, two proteins passed within-contrast BH adjustment for the midlobular-versus-endpoint-mean comparison (Fig. 4b,c). Hydroxyacid oxidase HAO1 and 3-hydroxyanthranilate 3,4-dioxygenase HAAO had mean paired differences of 0.656 and 0.651 on the log₂(*x* + 1) scale, with 95% confidence intervals of 0.631–0.680 and 0.600–0.703 and adjusted *P* values of 0.000386 and 0.00355, respectively. No protein met the within-contrast adjusted-*P* threshold in either individual-endpoint comparison. These candidates therefore describe the midlobular deviation from the average of both endpoints; the individual endpoint comparisons further resolve this spatial relationship.

Observed-only analysis refined the interpretation of these effects. The system used provenance-verified observation indicators to aggregate only original observations and repeat the comparisons on the same scale. HAO1 and HAAO retained positive effects relative to the endpoint mean, at 0.444 and 0.556, with 95% confidence intervals of 0.184–0.704 and 0.279–0.832. Compared with the full-matrix branch, their effects were smaller and intervals wider; both within-contrast BH-adjusted *P* values were 0.496. The branches thus preserved candidate direction while differing in precision and multiple-testing support. The three contrasts had 1,337, 1,454 and 1,437 testable proteins. In the midlobular-versus-central comparison, P17156 passed within-contrast BH adjustment (*q* = 0.0358), whereas its *q* value across the joint family of three contrasts was 0.1054 (Supplementary Fig. 5c). These results identify spatial candidates that can be investigated through mouse-level distributions, observation coverage and additional measurements.

This continuation separates a midlobular deviation from the endpoint mean from the two individual-endpoint differences. Reusing the matrix while changing the unit and contrasts produced paired estimates whose precision and observation support could be examined directly.

### Interpreting development and experimental context

Different research questions require evidence to be organized at different levels. Developmental studies ask whether proteins form coherent state-associated patterns; sample-handling studies connect technical context with measured differences; hematopoiesis studies link cell-state descriptions to candidate validation. Three complementary applications illustrate these uses, with offline analyses of saved outputs examining the conditions underlying interpretation (Fig. 5; Supplementary Figs. 6 and 7).

**Fig. 5.**
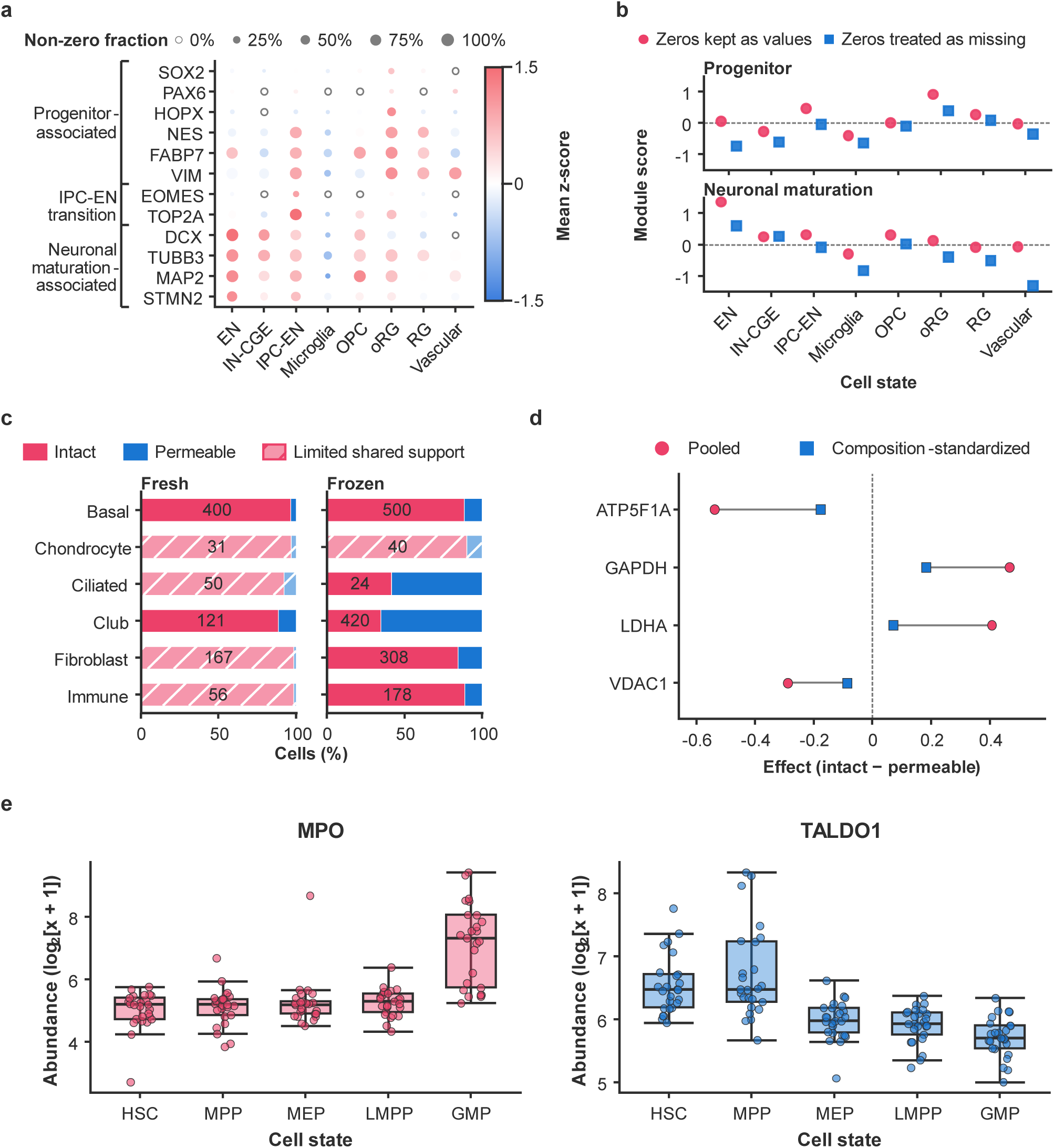
Interpreting developmental states and experimental context. **a**, Twelve marker proteins across eight labeled human brain states, grouped as progenitor, IPC-EN transition and neuronal maturation markers. Bubble area represents the nonzero fraction; an open symbol marks a zero fraction. Color indicates state-mean protein z scores, standardized across all 2,310 cells. Labels are available for 1,505 cells; 805 cells have no labels and are not shown as labeled states. State ordering does not represent an inferred trajectory. **b,** State-mean progenitor and neuronal maturation module scores under zero-retained and zero-as-missing conventions. Cell-level scores give equal weight to matched member genes before state-level averaging. Scores summarize relative protein abundance, not pathway activity. Nonzero-support analyses are shown in Supplementary Fig. 6. **c,** Membrane-state composition in 12 preservation-by-cell-type strata. Red and blue show intact and permeable fractions; numbers inside bars are total stratum cell counts. Hatching identifies limited common support, with fewer than five cells in either state. The other seven strata enter composition standardization. Complete counts are shown in Supplementary Fig. 7a. **d,** Pooled effects (circles) and composition-standardized effects (squares) for four fixed proteins, connected within each protein. Direction is intact minus permeable. Standardization uses the composition of intact cells within common-support strata. These are descriptive effects without estimated confidence intervals. **e,** Cell-level MPO and TALDO1 abundance on the log₂(*x* + 1) scale across five hematopoietic states, with 25 cells per state and 125 cells overall. The two facets display the same cells for two proteins, totaling 250 protein–cell values. Boxes show the 25th–75th percentiles, center lines the median and whiskers the most extreme observations within 1.5 interquartile ranges. Unit-level results from a different run and filtering set are shown in Supplementary Fig. 7b,c. RG, radial glia; oRG, outer radial glia; IPC-EN, intermediate progenitor-to-excitatory-neuron states; EN, excitatory neurons; HSC, hematopoietic stem cell; MPP, multipotent progenitor; MEP, megakaryocyte–erythroid progenitor; LMPP, lymphoid-primed multipotent progenitor; GMP, granulocyte–monocyte progenitor.

The developing human brain dataset addresses cell identity, neurogenesis and relationships between protein and transcript measurements^27^. Using existing labels for radial glia (RG), outer radial glia (oRG), intermediate progenitor-to-excitatory-neuron states (IPC-EN) and excitatory neurons (EN), the analysis examined protein programs distinguishing progenitor and neuronal states. Functional modules summarize the relative abundance of related proteins to describe their distribution across states. A progenitor module containing VIM and NES was relatively high in oRG, whereas a neuronal maturation module containing MAP2 and TUBB3 was relatively high in EN. These complementary patterns align with the neurodevelopmental context (Fig. 5a,b).

Such summaries need to be interpreted with member matching and nonzero-value support. Because the meaning of zeros in the input matrix was uncertain, offline analysis either retained zeros or treated them as missing. The relatively high patterns of both focal modules persisted, while score magnitudes changed with the convention (Fig. 5b; Supplementary Fig. 6). The module results summarize progenitor and neuronal states through the relative abundance of their constituent proteins, supporting the selection of state markers and subsequent measurement targets.

The protein leakage study asks how tissue handling affects measured cellular proteomes. The source study compared fresh and frozen mouse airway cells and used membrane permeability measurements to investigate sample integrity^28^. ScProteoAgent organized membrane-state comparisons, candidates and functional results in a matrix of 2,295 cells. Offline stratification of saved outputs then placed these differences within preservation and cell-type contexts. Of 12 preservation-by-cell-type strata, seven contained sufficient cells of both membrane states (Fig. 5c; Supplementary Fig. 7a). After standardization to the composition of membrane-intact cells within the common-support strata, ATP5F1A, GAPDH, LDHA and VDAC1 retained their effect directions, with magnitudes of 32.9%, 39.2%, 17.5% and 29.9% of the unstratified pooled effects, respectively (Fig. 5d). Intact-minus-permeable effects were positive for GAPDH and LDHA and negative for ATP5F1A and VDAC1. Their reduced magnitudes within matched preservation and cell-type contexts show why interpretation needs to consider both membrane-state associations and sample composition. The standardized and pooled estimates refer to different population weightings and support sets; their magnitude ratio describes the change between these summaries.

The early human hematopoiesis study combined single-cell proteomics and transcriptomics to examine stem and progenitor differentiation and validate selected protein functions^29^. In the existing cell-level analysis of the protein matrix, granulocyte–monocyte progenitors (GMPs) showed higher MPO and lower TALDO1 than hematopoietic stem cells (HSCs) (Fig. 5e). The higher TALDO1 abundance in HSCs relates to the source study’s investigation of this candidate. These cell-level directions connect the saved analysis with a candidate investigated in the source study.

Saved products also support examining these candidates at the experimental-unit level. In a separate run, none of the proteins in six estimable contrasts passed the BH threshold (Supplementary Fig. 7b,c), focusing the next question on whether candidate directions and magnitudes persist across more independent units. Cell-level results organize candidates, while unit-level results guide cross-unit validation.

These applications illustrate functional summarization, interpretation of sample context and organization of differentiation candidates. Across these settings, the interpretation depends on the level of aggregation, processing convention and experimental context.

## Discussion

ScProteoAgent organizes established proteomic calculations around research questions and preserves the context needed to continue an analysis. Its contribution combines a domain-specific computational system, a benchmark of connected research requirements and applications that relate statistical outputs to biological interpretation. Question-centric analysis defines the objects and comparisons; persistent analytical state retains their design and processing context; continuable scientific analysis makes these products available for the computations required by a later question. Together, these designs connect the result of one analysis to the inputs and decisions of the next.

Preserving experimental design and processing context connects established calculations to evolving questions. Processing frameworks provide standardized organization, quality control and interpretable modeling^12, 13^, while scientific agents coordinate tools and support iterative exploration^20, 23, 24, 25^. ScProteoAgent brings these capabilities into a domain-specific design: groups, contrast directions, experimental units and matrix state enter tool parameters and remain associated with results. In the liver case, a researcher supplied mouse identifiers and spatial questions, and the system reused the matrix while aggregating observations and calculating new paired contrasts. Reuse therefore involved retaining valid inputs and performing the newly required computation. This example makes the role of analytical state concrete without requiring the original statistical methods to be replaced.

The benchmark evaluates this framework against real analytical requirements. Progressive tasks connect grouping, candidates and integrated interpretation within a study, while complementary experimental conditions provide different analytical settings. Content requirements supported by source studies and a fixed six-dimensional rule function establish a repeatable basis for comparison. The evaluation addresses complete research outputs through content coverage, quantitative evidence, interpretation and artifact records. ScProteoAgent achieved the highest composite score in all 11 studies, with advantages concentrated in research content, matrix evidence and the organization of interpretation and reports. These scores characterize coverage and reporting in archived outputs under historical generation conditions. The benchmark also preserves tasks, source studies and evaluation rules for future comparisons.

The two principal cases demonstrate complementary research uses. PiSPA connects migration phenotypes, population composition and cluster-associated candidates. Original-observation records distinguish well-supported measurements from differences sensitive to imputation, and the GO background comparison identifies which functional interpretations depend on the reference population. DVP uses saved outputs to address a more specific spatial question: mouse-level contrasts distinguish a midlobular deviation from the endpoint mean from differences relative to individual endpoints. Retaining full-matrix and observed-only branches exposes the associated changes in precision and multiple-testing support. Developmental, membrane-state and hematopoietic applications extend this approach to functional summaries, sample composition and replicate structure. These analyses organize existing biological relationships into candidates and comparisons that can guide subsequent measurement.

Persistent analytical state also provides a direction for public-data reuse. Single-cell proteomic studies accumulate information on cell states, spatial locations and experimental responses; the same matrix can address questions posed later. Keeping data, metadata and processing records findable, understandable and reusable underpins the long-term value of public resources^16, 30^. quantms demonstrates scalable reanalysis of public proteomic data^31^, while RO-Crate and its workflow-run extensions organize research products, computational provenance and object relationships^18, 19^. These approaches provide complementary foundations for reuse. ScProteoAgent uses relationships among questions, designs and results during research interaction, enabling researchers to locate applicable inputs and translate new questions into computation. Connections to standardized data frameworks could support interactive investigation of public data, in which researchers progressively ask questions, obtain results with explicit computational support and preserve updated state for subsequent exploration.

Further development should extend the demonstrated continuation path to more tasks and longer interactions. Dependency-aware recalculation after a question changes, context selection and interpretation are tractable targets for systematic evaluation. Knowledge connections with explicit species and protein identity could further support the conversion of matrix-derived candidates into testable biological hypotheses. Advances linking proteomic measurements to imaging, spatial location and functional phenotypes will provide richer experimental context^5, 6^. The system, benchmark and applications developed here provide a foundation for directing growing single-cell proteomic resources toward specific, testable questions that can be pursued through subsequent analysis.

## Methods

### Study design and benchmark

We analyzed published protein matrices and sample information from 11 studies, comprising 5,262 measured objects. Migration and spatial data derive from PiSPA and liver zonation studies^5, 26^. Developmental datasets cover induced pluripotent stem cells, human brain and early hematopoiesis^27, 29, 32^. Other studies address immune and neuronal states^33, 34, 35, 36^, protein turnover and membrane integrity^28, 37^. Acquisition, identification and primary quantification were performed in the source studies.

Each benchmark instance comprises a protein matrix, sample metadata, natural-language requirements, source-supported reference conclusions and scoring criteria. We manually organized five requirements and three reference conclusions per study, yielding 55 tasks and 33 reference conclusions. The five requirements were supplied together and progress from grouping or state identification to candidates, functional interpretation and integrated judgments. Matrices and metadata were aligned by sample identifiers, with annotation columns separated from quantitative values. Observation counts refer to cells, spatial sampling units or low-input samples, as appropriate, rather than uniformly to independent biological replicates.

Cross-system evaluation uses 88 archived outputs. Application figures use designated runs and offline analyses of their saved products; the liver follow-up uses the continuation interface.

### Agent implementation and analytical state

ScProteoAgent uses a Python/LangGraph state graph with Planner, Executor, Analyzer, Reporter, Critic and Backtrack nodes (Fig. 1a). Shared state stores the task, plan, current step, tool returns, interim summaries and report segments. The model plans tool use and interpretation; registered computational tools perform numerical operations and return structured summaries and artifact locations. Report assembly combines generated text with deterministic result tables.

The analysis design records sample and protein identifiers, gene annotations, groups, target and reference directions, experimental units, species and matrix-processing state. Target minus reference defines effect direction. Research intent, analysis design, data state and result state are retained through shared objects and saved design files, outputs and artifact indexes. Evidence records link reported findings to statistical products and their sources.

For continuation, the interface reads the parent task, design and artifact index, including matrix identity, processing state and dependencies. Parent artifacts remain read-only; an updated design determines any new aggregation or testing, and new outputs are saved separately. The historical comparison and subsequent liver continuation use their respective implementations. GPT-5.5 was the base model for ScProteoAgent in the principal comparison; case-specific configurations were recorded during analysis.

### Matrix processing and differential analysis

Global filtering applies a nonmissing-fraction threshold across all samples; group-wise filtering requires the threshold in each specified group. Where used, half-minimum imputation replaces missing entries with half the smallest observed value for that protein on the current scale. Logarithmic transformation uses log_2_(*x* + 1) ; previously processed inputs retain their recorded scales.

The tool library includes R/limma^38^ and Python statistical backends. The initial HeLa, liver and specified cell-level hematopoietic analyses used two-sided Welch *t*-tests, with group mean differences on the analysis scale. Benjamini–Hochberg (BH) adjustment was applied within each contrast^39^. Initial PiSPA and liver candidate selection required adjusted *P* ≤ 0.05 and absolute effect ≥ 0.25. The mouse-level liver analyses used the tests and correction families specified below, without this additional effect cutoff. ComBat is available^40^, but no new ComBat correction was applied in these Python case branches.

### Functional analysis and GO backgrounds

Functional modules use saved member lists matched to matrix gene annotations. Module scores summarize member abundance; the brain-specific standardization and weighting procedures are described below. The tool library also supports rank-based enrichment using GSEA^41^ and external annotation interfaces.

HeLa GO analysis used human biological-process, cellular-component and molecular-function sets derived from MSigDB v2026.1.Hs. Their union contained 19,591 gene symbols. For each cluster contrast, higher- and lower-abundance queries required adjusted *P* ≤ 0.05 and absolute mean difference ≥ 0.25. Annotated gene symbols were split, deduplicated and restricted to the annotation universe. One-sided hypergeometric tests evaluated overlaps. The three namespaces were pooled, and BH adjustment included every term with at least one query-gene overlap within each contrast and direction. Zero-overlap terms were excluded. Query and testing-family sizes are listed in Supplementary Methods.

An offline sensitivity analysis substituted the 4,227 annotated genes represented in testable differential-table rows as the experimental background. All 4,688 rows had finite effects, *P* values and adjusted *P* values; one lacked annotation. The background was identical across contrasts and contained all annotation-matched query genes. Background and term sizes changed, while query membership, overlaps and correction families remained fixed. Complete results were retained before applying *q* ≤ 0.05. Historical calculations reproduced all 90 saved records before interpreting the alternative background.

Fig. 3e retains eight terms selected as the four smallest adjusted *P* values per direction in C1 versus C3. Its historical source tables saved up to 15 significant terms per query, so absence from those tables does not establish nonsignificance. Both backgrounds were evaluated at all 24 fixed term–contrast positions.

### Archived-output evaluation and paired comparisons

ScProteoAgent, Biomni, SpatialAgent, BioAgent, CellVoyager, Codex, Claude Code and Hermes each contributed outputs for 11 studies. Scores were deterministically reproduced with the frozen original six-dimensional program, using reports and their archived run directories. Reference texts, report mappings and archive paths were fixed. Historical language-model adjustments and additional normalization were excluded.

The 100-point total comprises research content (35), matrix evidence (20), interpretive boundaries (15), artifacts and run-state records (15), Chinese report structure and expression (10), and points retained after penalties (5). The same program, weights and input conventions apply to all systems. Dimension scores are rounded to two decimal places before aggregation. Supplementary Methods describes the matching rules.

The evaluation measures archived-output compliance with research-content and delivery requirements; case statistics are checked separately. Evaluation-requirement files occur in 33 of 88 run directories, and reference-candidate names occur in 59 reports. The archived THP-1 case links reference-derived requirements to candidate and module tables in the assembled report. Their use in model prompts remains unresolved. These historical conditions define an output comparison, not a controlled comparison with reference material held out.

System means use the 11 study-level totals. Paired differences are ScProteoAgent minus each comparator, with studies as resampling units. Percentile-bootstrap 95% confidence intervals use 20,000 resamples and one shared (20,000 × 11) index matrix from NumPy Generator/PCG64 (seed 20260919). Endpoints are the 2.5th and 97.5th percentiles, calculated by linear interpolation.

### HeLa migration analysis

The input contained 5,191 protein rows and 89 cells with source-provided cluster and migration labels. Requiring at least 10% nonmissing values separately in migrated and control cells retained 4,688 proteins. Half-minimum imputation preceded log_2_(*x* + 1) transformation, without batch correction. For three-component PCA, proteins were centered and scaled to unit variance across cells. Cluster comparisons used two-sided Welch tests with separate BH families.

The offline migration-label comparison used the same matrix, with 46 migrated and 43 control cells and one 4,688-protein BH family. Fig. 3d shows *q* < 0.05 without the cluster-candidate effect cutoff. The composition odds ratio compares C1 with C2: (35 × 26)/(7 × 5) = 26.0.

Original and imputed TBC1D10B values were identified by entry-level matching to the pre-imputation matrix. Of 89 values, 31 were imputed: 10/42 in C1, 7/31 in C2 and 14/16 in C3. Box plots summarize all processed values, with observation identities displayed separately (Supplementary Fig. 4).

### Liver zonation and continued analysis

The liver input comprised 2,149 protein rows and 45 sampling sites. Global filtering at 10% nonmissing values retained 1,859 proteins, followed by half-minimum imputation and log_2_(*x* + 1) transformation. Initial sampling-site comparisons used Welch tests and within-contrast BH adjustment for central–midlobular, central–portal and midlobular–portal contrasts. For continuation, the researcher supplied mouse identifiers and three spatial questions. Sample identifiers linked the 45 sites to five mice, three zones per mouse and three sites per zone. The agent updated the design and averaged transformed abundance within each mouse and zone, yielding 15 units. Identifier and zone mappings were checked for duplicates, unmatched samples and conflicts.

For protein *p* and mouse *u*, let 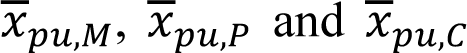 denote midlobular, portal and central means. The contrasts were

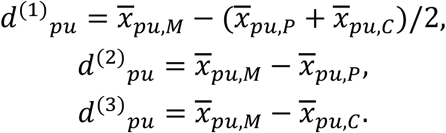

Effects are mean within-mouse differences among mice with finite values in all required zones. Two-sided one-sample *t*-tests compare these differences with zero, requiring at least three valid pairs. Insufficient pairing or zero variance produces no valid *P* value. Unadjusted 95% confidence intervals are 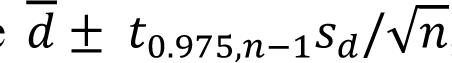; paired counts are reported for each protein. The full-matrix branch includes imputed values. The observed-only branch masks those entries before aggregation, leaving units without original observations missing. Both branches use the same locked matrix, scale and contrasts. BH adjustment is performed separately within each contrast and across all three contrasts within each branch (*q* < 0.05). Joint families contain 5,576 finite *P* values among 5,577 full-matrix rows and 4,228 observed-only values. Family-specific counts are given in Supplementary Fig. 5. Aggregates, effects, *P* values and paired counts were checked independently.

### Brain modules and zero-value sensitivity

The brain matrix contained 4,279 protein rows and 2,310 cells. Protein rows were standardized across cells using the sample standard deviation, 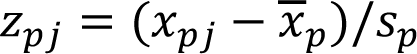; zero-variance rows were set to missing. Original module scores averaged matched member-row z scores within each cell, ignoring missing values. Multiple protein rows for one gene contributed separately. Zero-value sensitivity either retained zeros or treated exact zeros as missing before recalculating means, standard deviations and scores. Fig. 5b uses equal-gene weighting: matched row z scores are first averaged within genes, then across member genes within each cell. Cell scores are averaged by state. Offline Spearman correlations relate state-mean module scores to nonzero fractions, using either nine groups including the unlabeled pool or eight labeled states. Nonzero fractions describe matrix support; the biological meaning of zeros is unresolved.

### Composition standardization in airway cells

The normalized input contained 1,223 proteins and 2,295 cells, including 1,838 intact and 457 permeable cells. Values could be positive or negative. All proteins were complete and passed the global 90% nonmissing threshold; no imputation or additional log transformation was applied. Preservation condition and cell type defined 12 strata. Seven contained at least five cells in each membrane state and formed the common-support set *S*.

For protein *p* in stratum *s*, *d_ps_* is the intact-minus-permeable mean difference. Weights follow intact-cell composition, *w_s_* = *n_s_*_,Intact_/1838, and are renormalized over *S*:

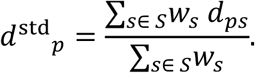

The pooled effect is the unstratified difference across all cells. For four fixed candidates, the magnitude ratio is |*d*^std^*_p_*|/|*d*^pooled^*_p_*| , with direction reported separately. This offline descriptive comparison changes population weighting and support; it does not estimate a causal contribution of leakage.

### Hematopoietic cell- and unit-level analyses

Cell-level analysis used 125 cells, with 25 each from HSC, MPP, MEP, LMPP and GMP states. Global 95% nonmissing filtering retained 336 proteins. Half-minimum imputation and log_2_(*x* + 1) transformation preceded GMP-minus-HSC Welch testing and within-contrast BH adjustment.

A separate run applied 10% global filtering and retained 1,550 proteins for unit-level analysis. Values were averaged by state within metadata donor labels, excluding mixed-source labels according to that run’s rules. Paired *t*-tests required at least three labels containing both states. Six contrasts had three to five pairs, with BH adjustment within each 1,550-protein family; four contrasts with two pairs were not tested. The runs differ in filtering, aggregation and testing, so their comparison does not isolate an experimental-unit effect.

### Software and result records

Python/LangGraph coordinates the workflow; NumPy^42^, pandas and SciPy^43^ support matrix operations and statistical testing. R packages provide additional tool backends. Run-specific model identities, environments and parameters were recorded during analysis.

## Acknowledgements

This work was supported by the New Generation Artificial Intelligence–National Science and Technology Major Project (2025ZD0122805, to Y.W. and K.D.), the National Natural Science Foundation of China (22404145, to Y.W., and 62676353, to K.D.), the Zhejiang Provincial Key Research and Development Program (2026C02A1108, to Y.W.) and the Natural Science Foundation of Zhejiang Province (LMS26B050004, to Y.W.).

## Author contributions

R.H., Y.W., and K.D. conceived the project. R.H. and Y.W. performed the majority of the coding work, including Agent development and implementation of inference workflows. Y.W. and K.D. provided guidance on model design and optimization. Y.W. and W.W. performed dataset curation, post-inference analyses, and visualization. S.W. and K.D. provided the computational resources. R.H., Y.W., and K.D. wrote the initial draft. P.C. and Z.Z. contributed visualizations. Y.W., K.D., and S.W. supervised the project. All authors revised the paper.

