## Supplementary information for "ScProteoAgent enables natural-language-driven single-cell proteomics analysis and interpretation"

### Contents

Supplementary Figs. 1–7

Supplementary Methods: additional scoring details and GO query specifications

Supplementary References

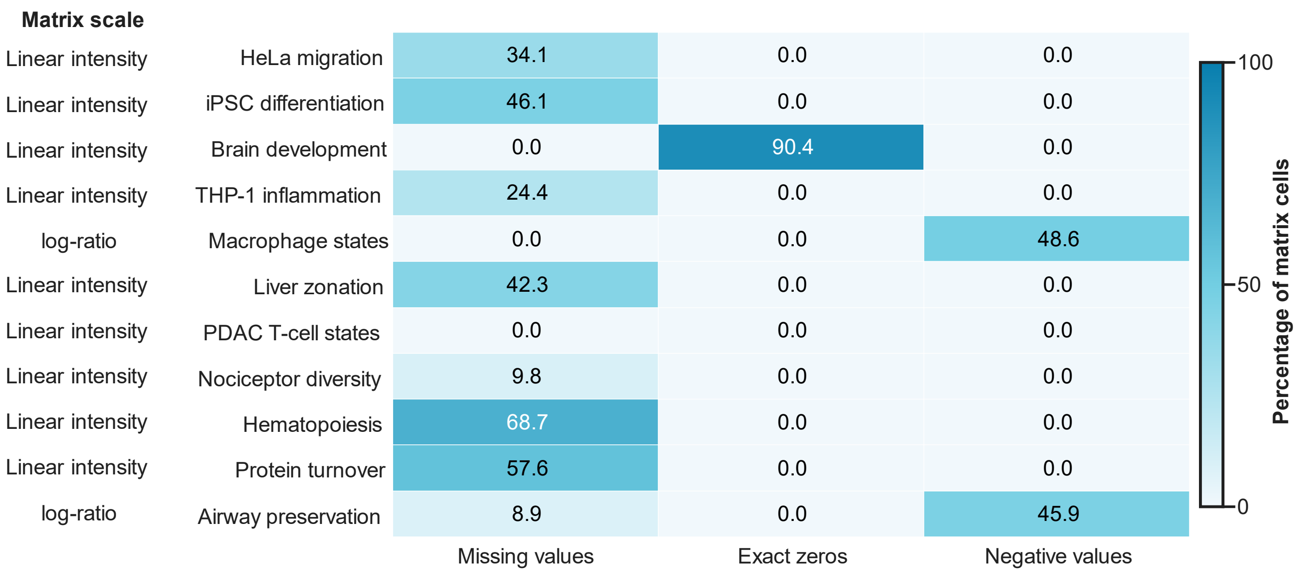


**Supplementary Fig. 1 | Numerical properties of the input matrices.**

Fractions of missing, zero and negative entries in the 11 input matrices, using all matrix entries as the denominator. Cell values are percentages; the color scale spans 0–100. Recorded input scales are indicated by study. Categories are mutually exclusive but omit positive entries, so displayed percentages need not sum to 100%. Zero values are not assigned a common biological meaning.

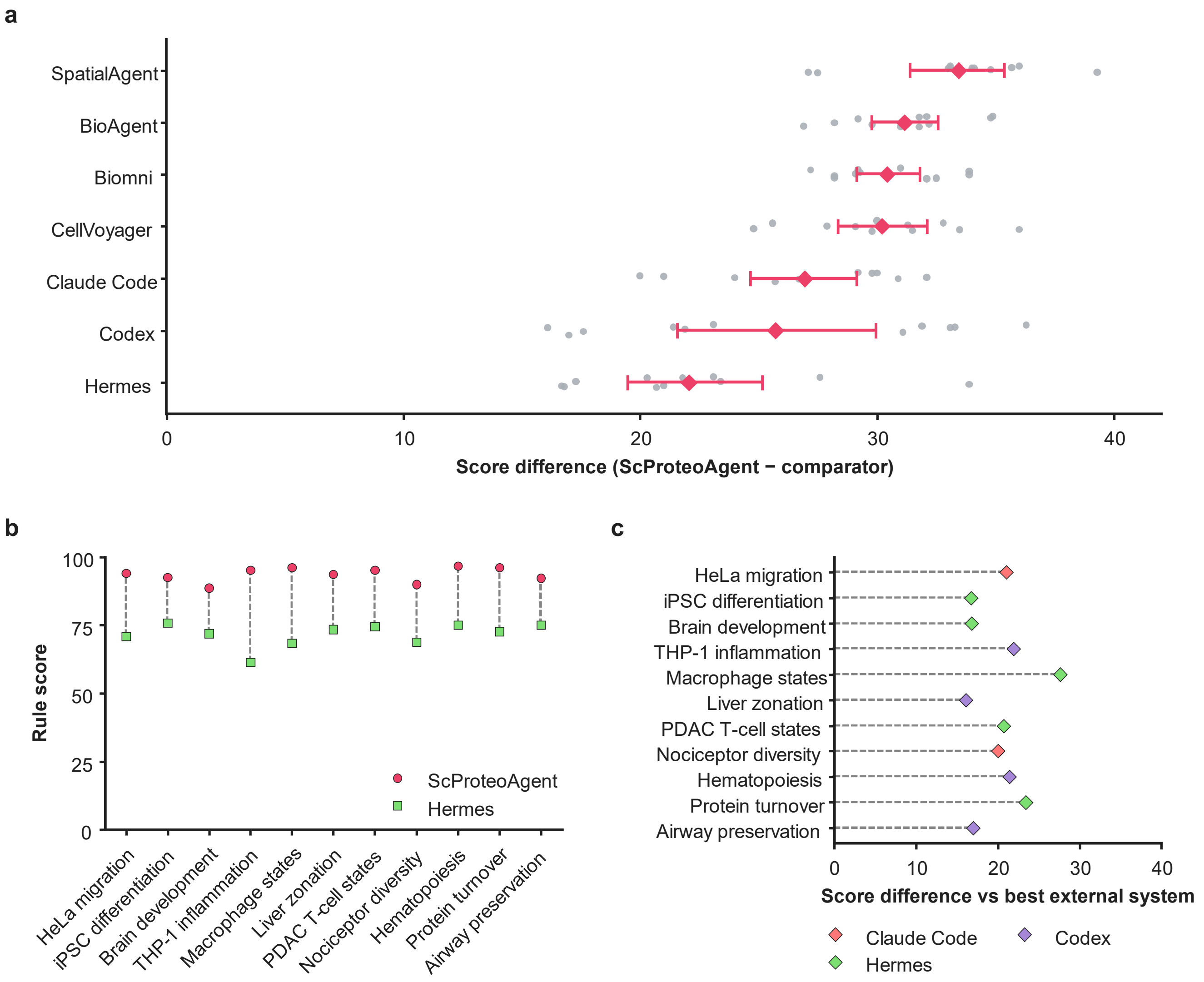


**Supplementary Fig. 2 | Paired study-level score comparisons.**

**a,** ScProteoAgent-minus-comparator rule scores for seven systems across 11 studies, giving 77 paired differences. Gray points are study-level differences, red filled diamonds are mean differences and horizontal lines are percentile-bootstrap 95% confidence intervals from 20,000 study-level resamples, with one index matrix shared across comparisons (Methods). **b,** ScProteoAgent and Hermes, the comparator with the highest mean score, for each study. Dashed lines connect the two scores within a study, yielding 22 points and 11 pairs. **c,** The difference between ScProteoAgent and the highest-scoring external system within each study. Diamonds mark differences and dashed lines connect zero to each estimate. Color identifies the corresponding external system, which can differ among studies. All panels use the same 88 outputs and six-dimensional rule evaluation.

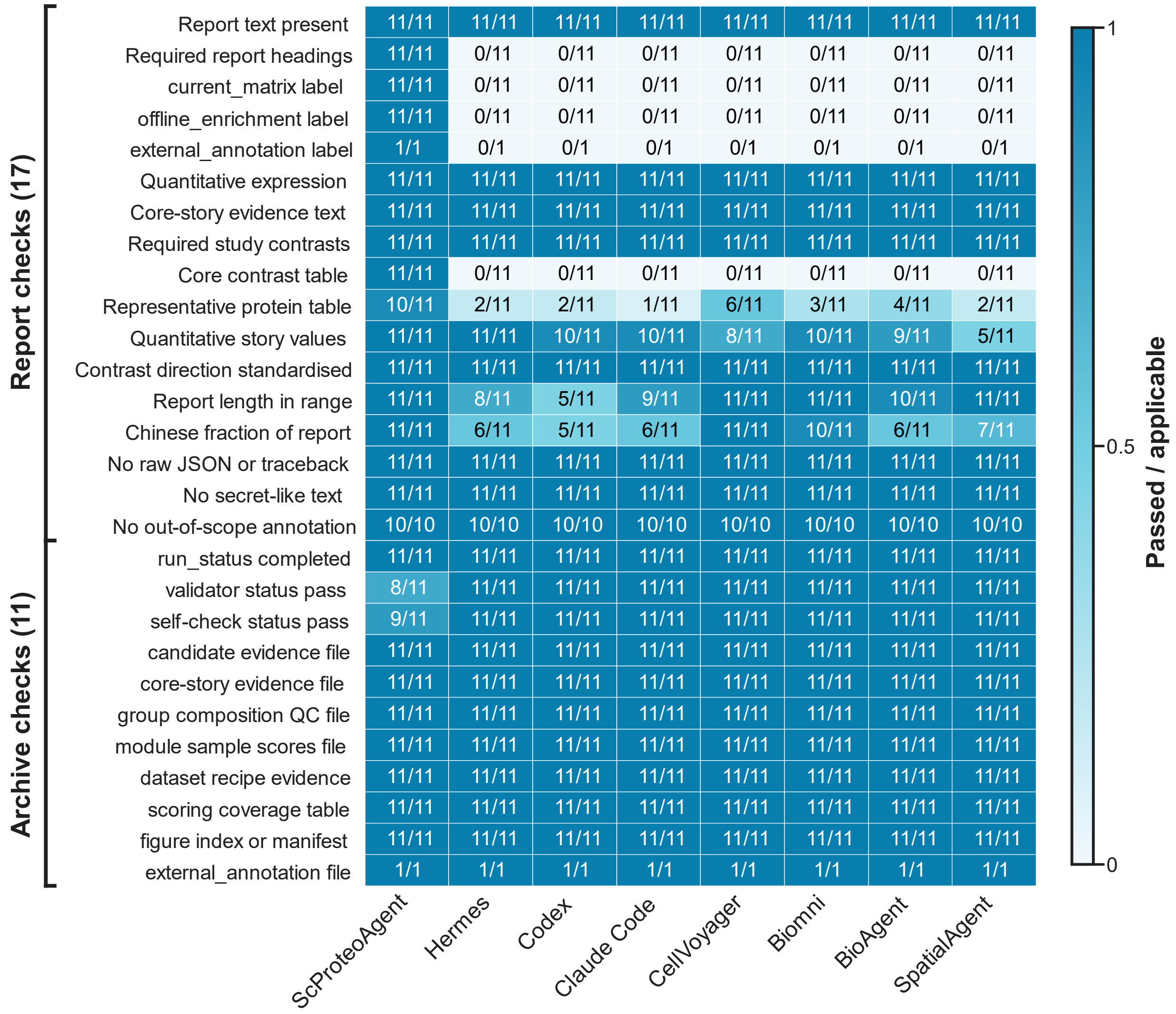
Report checks (17)
Archive checks (11)
Report text present 11/11 11/11 11/11 11/11 11/11 11/11 11/11 11/11
Required report headings 11/11 0/11 0/11 0/11 0/11 0/11 0/11 0/11
current_matrix label 11/11 0/11 0/11 0/11 0/11 0/11 0/11 0/11
offline_enrichment label 11/11 0/11 0/11 0/11 0/11 0/11 0/11 0/11
external_annotation label 1/1 0/1 0/1 0/1 0/1 0/1 0/1 0/1
Quantitative expression 11/11 11/11 11/11 11/11 11/11 11/11 11/11 11/11
Core-story evidence text 11/11 11/11 11/11 11/11 11/11 11/11 11/11 11/11
Required study contrasts 11/11 11/11 11/11 11/11 11/11 11/11 11/11 11/11
Core contrast table 11/11 0/11 0/11 0/11 0/11 0/11 0/11 0/11
Representative protein table 10/11 2/11 2/11 1/11 6/11 3/11 4/11 2/11
Quantitative story values 11/11 11/11 10/11 10/11 8/11 10/11 9/11 5/11
Contrast direction standardised 11/11 11/11 11/11 11/11 11/11 11/11 11/11 11/11
Report length in range 11/11 8/11 5/11 9/11 11/11 11/11 10/11 11/11
Chinese fraction of report 11/11 6/11 5/11 6/11 11/11 10/11 6/11 7/11
No raw JSON or traceback 11/11 11/11 11/11 11/11 11/11 11/11 11/11 11/11
No secret-like text 11/11 11/11 11/11 11/11 11/11 11/11 11/11 11/11
No out-of-scope annotation 10/10 10/10 10/10 10/10 10/10 10/10 10/10 10/10
run_status completed 11/11 11/11 11/11 11/11 11/11 11/11 11/11 11/11
validator status pass 8/11 11/11 11/11 11/11 11/11 11/11 11/11 11/11
self-check status pass 9/11 11/11 11/11 11/11 11/11 11/11 11/11 11/11
candidate evidence file 11/11 11/11 11/11 11/11 11/11 11/11 11/11 11/11
core-story evidence file 11/11 11/11 11/11 11/11 11/11 11/11 11/11 11/11
group composition QC file 11/11 11/11 11/11 11/11 11/11 11/11 11/11 11/11
module sample scores file 11/11 11/11 11/11 11/11 11/11 11/11 11/11 11/11
dataset recipe evidence 11/11 11/11 11/11 11/11 11/11 11/11 11/11 11/11
scoring coverage table 11/11 11/11 11/11 11/11 11/11 11/11 11/11 11/11
figure index or manifest 11/11 11/11 11/11 11/11 11/11 11/11 11/11 11/11
external_annotation file 1/1 1/1 1/1 1/1 1/1 1/1 1/1 1/1
ScProteoAgent
Hermes
Codex
Claude Code
CellVoyager
Biomni
BioAgent
SpatialAgent
Passed / applicable
1
0.5
0

**Supplementary Fig. 3 | Pass fractions for fixed rule checks.**

Eight systems evaluated on 28 fixed checks, comprising 17 report checks and 11 archive checks. Color encodes passing divided by applicable objects; both numerator and denominator are shown. Applicable denominators are 1, 10 or 11 according to the check and evaluated objects. Nonapplicable objects are excluded from the denominator. Term, structure and artifact-presence checks measure rule compliance of archived outputs and are distinct from verification of numerical values and scientific interpretation.

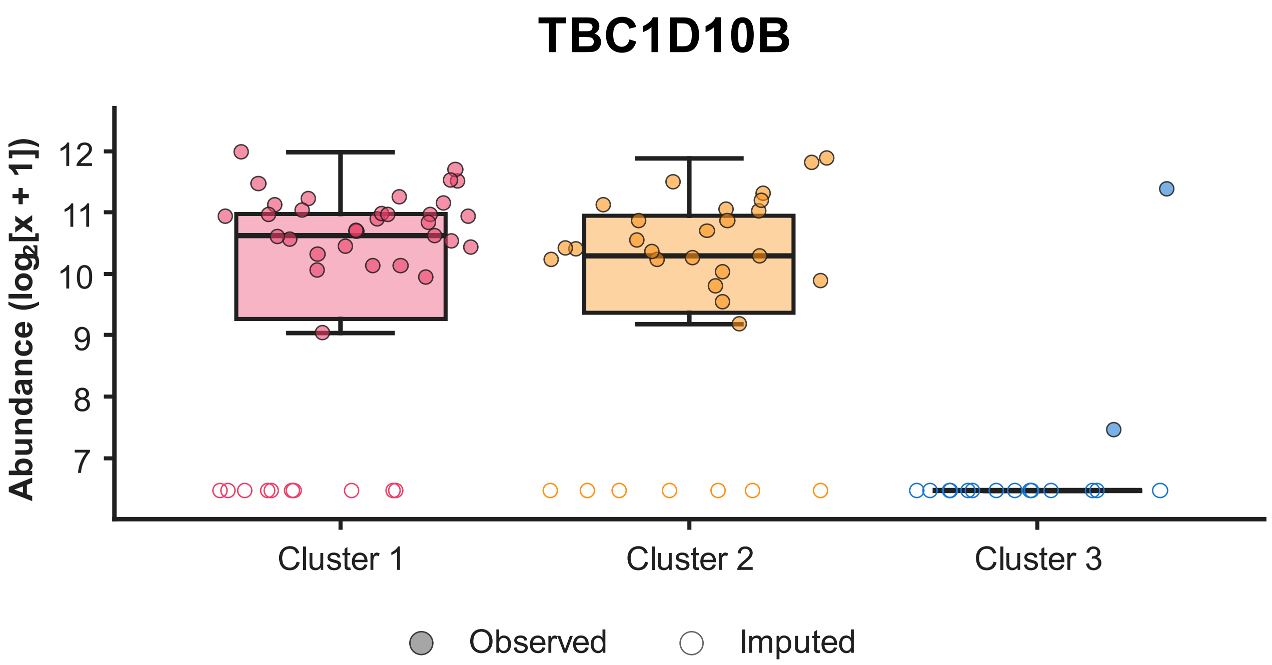


**Supplementary Fig. 4 | Observation support for the cluster-associated candidate TBC1D10B.** Processed TBC1D10B abundance for 89 HeLa cells, grouped by the three clusters and shown on the $\log_2(x + 1)$ scale. Entry-level matching to the pre-imputation matrix identifies original observations (filled points) and imputed values (open points). There are 31 imputed values among 89 entries: 10/42 in C1, 7/31 in C2 and 14/16 in C3. C3 therefore contains two original observations. Boxes summarize all processed values within a cluster, with the 25th–75th percentiles, median and whiskers extending to the most extreme observations within 1.5 interquartile ranges. Observation identities support interpretation of Fig. 3c.

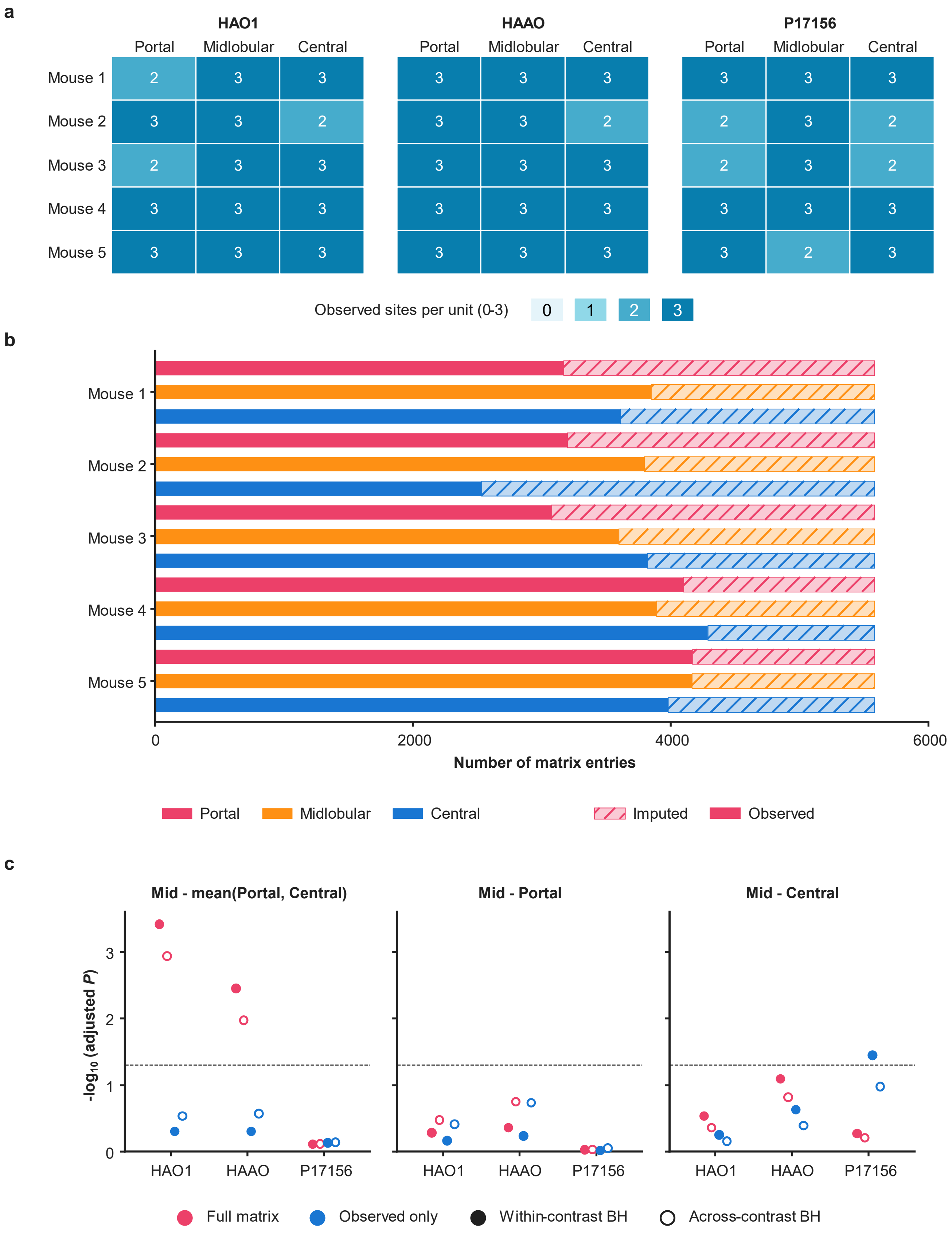
a
HAO1
HAAO
P17156
Portal
Midlobular
Central
Mouse 1
Mouse 2
Mouse 3
Mouse 4
Mouse 5
Observed sites per unit (0-3)
0
1
2
3
b
0
2000
4000
6000
Number of matrix entries
Portal
Midlobular
Central
Imputed
Observed
c
Mid - mean(Portal, Central)
Mid - Portal
Mid - Central
-log10 (adjusted P)
HAO1
HAAO
P17156
Full matrix
Observed only
Within-contrast BH
Across-contrast BH

**Supplementary Fig. 5 | Observation support and correction families in liver zonation.**

**a,** Original observation counts for three candidates in 15 mouse-by-zone units, with three sampling sites per unit. Cell values and color indicate observed sites out of three. **b,** Observed and imputed matrix entries by mouse and zone. Rose, orange and blue identify portal, midlobular and central zones. Unhatched segments represent observations; hatched segments represent imputed entries. Each unit contains 1,859 × 3 = 5,577 protein–site entries. **c,** BH-adjusted $P$ values for 18 candidate estimates, each evaluated within its contrast and across the three contrasts, yielding 36 values. Rose and blue distinguish full-matrix and observed-only branches. Filled symbols denote within-contrast BH; open symbols denote across-contrast BH. Both adjustments are performed separately within each branch. The dashed line marks $q = 0.05$. The three within-contrast families contain 1,859/1,858/1,859 finite $P$ values in the full-matrix branch and 1,337/1,454/1,437 in the observed-only branch. Joint families therefore contain 5,576 and 4,228 values, respectively. One full-matrix result row lacks a valid $P$ value. Observed-only P17156 in the midlobular–central contrast has within-contrast $q = 0.0358$ and across-contrast $q = 0.1054$. Corresponding confidence intervals in Fig. 4c are unadjusted for multiplicity.

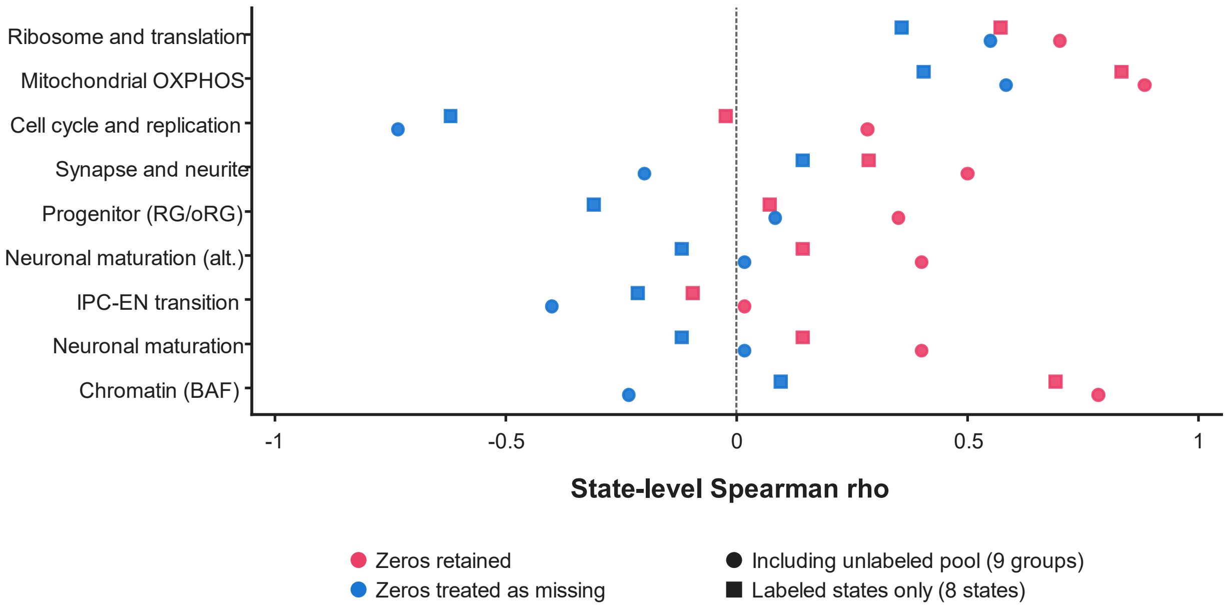


**Supplementary Fig. 6 | Module scores and nonzero-value support.**

Spearman correlations between state-mean module scores and nonzero fractions for nine modules. Rose denotes zeros retained; blue denotes zeros treated as missing. Circles include nine groups, comprising eight labeled states and the unlabeled pool; squares include only the eight labeled states. The plot contains 36 coefficients calculated from group summaries. The dashed line marks zero correlation. Nonzero fractions describe numerical matrix support, rather than confirmed detection rates.

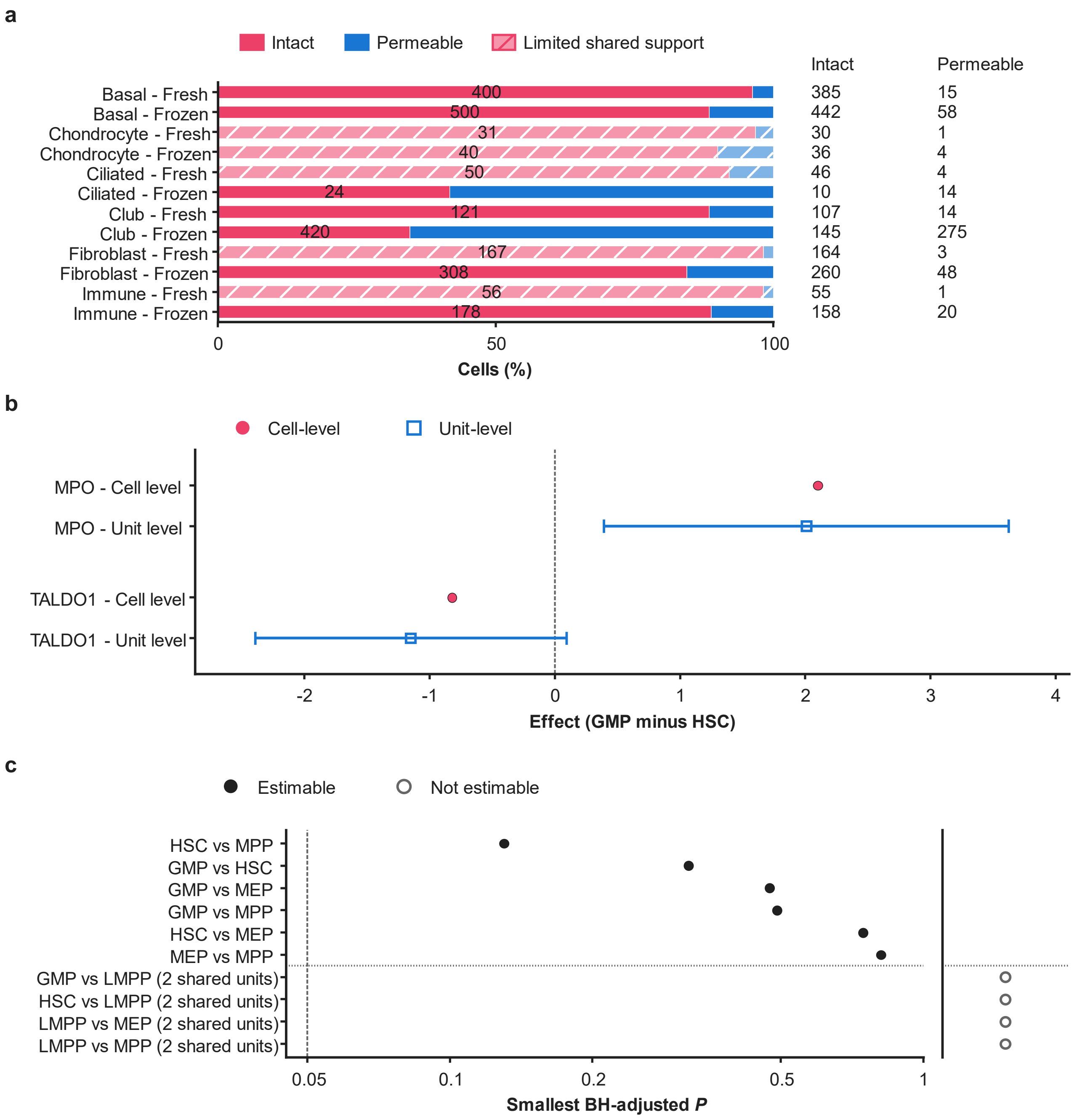
a
Intact
Permeable
Limited shared support
Intact
Permeable
Basal - Fresh
Basal - Frozen
Chondrocyte - Fresh
Chondrocyte - Frozen
Ciliated - Fresh
Ciliated - Frozen
Club - Fresh
Club - Frozen
Fibroblast - Fresh
Fibroblast - Frozen
Immune - Fresh
Immune - Frozen
400
500
31
40
50
24
121
420
167
308
56
178
385 15
442 58
30 1
36 4
46 4
10 14
107 14
145 275
164 3
260 48
55 1
158 20
0
50
100
Cells (%)
b
Cell-level
Unit-level
MPO - Cell level
MPO - Unit level
TALDO1 - Cell level
TALDO1 - Unit level
-2
-1
0
1
2
3
4
Effect (GMP minus HSC)
c
Estimable
Not estimable
HSC vs MPP
GMP vs HSC
GMP vs MEP
GMP vs MPP
HSC vs MEP
MEP vs MPP
GMP vs LMPP (2 shared units)
HSC vs LMPP (2 shared units)
LMPP vs MEP (2 shared units)
LMPP vs MPP (2 shared units)
0.05
0.1
0.2
0.5
1
Smallest BH-adjusted P

**Supplementary Fig. 7 | Preservation context and hematopoietic unit-level results.**

**a,** Membrane-state composition within preservation-by-cell-type strata. Red and blue denote intact and permeable fractions. Bar labels show stratum totals and adjacent values show counts for each state. Hatching identifies strata with fewer than five cells in either state. This is a detailed count view of the same 2,295 cells shown in Fig. 5c. **b,** Cell-level and unit-level effects for MPO and TALDO1, oriented as GMP minus HSC. Effects retain the scales and definitions of their respective analyses; positive values indicate higher abundance in GMP. The cell-level analysis estimates a mean difference after $\log_2(x + 1)$ transformation, as specified in Methods. Circles show cell-level estimates, open squares unit-level estimates, and intervals are the saved unit-level 95% confidence intervals. The estimates come from different runs retaining 336 and 1,550 proteins, with different aggregation units and testing families. Differences between these estimates do not isolate an experimental-unit effect. Unit-level BH-adjusted $P$ values are 0.7507 for MPO and 0.7626 for TALDO1; neither passes 0.05 within its testing family. **c,** Estimability of ten predefined unit-level contrasts. Six have three to five shared units and test 1,550 proteins per contrast. Four have only two shared units and are represented by open symbols without assigned $P$ values. Filled points show the smallest BH-adjusted $P$ value in each estimable contrast. The horizontal axis is logarithmic and the dashed line indicates 0.05. None of the six estimable contrasts has a $q < 0.05$ protein. The smallest value is 0.1301 for HSC versus MPP; GMP versus HSC has a minimum of 0.3189. GMP, granulocyte–monocyte progenitor; HSC, hematopoietic stem cell; MPP, multipotent progenitor; BH, Benjamini–Hochberg.

### Supplementary Methods

#### Additional scoring details

The frozen scorer evaluates each report together with its archived run directory. Research-content checks combine reference-term coverage, required contrasts, key findings and core-story content. Terms are extracted from reference conclusions and scoring criteria, filtered with a fixed stop-word list, ordered by frequency and limited to 80 distinct terms. Coverage is the fraction matched in the report.

Matrix-evidence checks identify quantitative expressions, candidate evidence and result access. Boundary checks identify scope statements and external annotation records where applicable. Artifact checks cover result tables, module records, dataset-recipe information, figure inventories and run status. Report-language checks assess section organization, Chinese-character fraction, body length, core-table labels and raw diagnostics. Penalties address absent content, incomplete status, exposed diagnostics or suspected credentials, and specified contrast or annotation inconsistencies. Caps and clipping follow the frozen specification. Each dimension is rounded to two decimal places before the rounded total is calculated; system means use the 11 study totals. Supplementary Fig. 3 uses each check's applicable denominator. Dimension weights and paired-bootstrap procedures are specified in Methods.

#### GO query and correction-family sizes

Higher- and lower-abundance queries are defined relative to the first-named cluster. BH adjustment[1] pools the three GO namespaces within each contrast and direction, including only terms with at least one query-gene overlap. Query membership and correction families were identical under both backgrounds.

In the order C1–C2 higher/lower, C1–C3 higher/lower and C2–C3 higher/lower, query sizes were 303/1,910, 575/292 and 1,245/39 genes. Corresponding testing-family sizes were 4,829/7,759, 6,165/3,887 and 7,465/1,187 terms. Significant-term counts changed from 826/851, 874/276 and 822/16 under the historical background to 382/80, 144/19 and 32/0 under the experimental background.

Significance is $q \leq 0.05$. Backgrounds contain 19,591 and 4,227 genes, respectively. These complete-query counts differ from the fixed 24-position display comparison in the main text. Fig. 3e retains saved historical results; its 11 open squares indicate absence from the truncated saved tables. Reconstruction supplies statistics for all positions.
